# RNA Modifications and Prp24 Coordinate Lsm2-8 Binding Dynamics during *S. cerevisiae* U6 snRNP Assembly

**DOI:** 10.1101/2025.02.25.639938

**Authors:** Ye Liu, Yuichiro Nomura, Samuel E. Butcher, Aaron A. Hoskins

## Abstract

In eukaryotes, the process of intron removal from nuclear pre-mRNA is performed by the spliceosome, a dynamic molecular machine composed of small nuclear ribonucleoproteins (snRNPs; U1, U2, U4, U5, and U6) and dozens of other protein splicing factors. The U6 snRNP contains the U6 snRNA and the proteins Prp24 and Lsm2-8 heteroheptamer. A key feature of the snRNP is a modified U6 snRNA 3’ end, which in *S. cerevisiae* (yeast) contains a 3’ phosphate. U6 plays an essential role in splicing, and the U6 snRNP must be completely disassembled for splicing to occur. Once splicing is finished, the snRNP must then be reassembled to participate in a subsequent splicing reaction. While splicing efficiency depends on rapid U6 snRNP assembly, this process has not yet been kinetically characterized. Here, we use colocalization single molecule spectroscopy (CoSMoS) to dissect the kinetic pathways of yeast U6 snRNA association with the Lsm2-8 complex and their dependence on the Prp24 protein and post-transcriptional snRNA modification. In the absence of 3’ end processing, Lsm2-8 association with the RNA is highly dependent on Prp24. However, processed RNAs can rapidly recruit Lsm2-8 in Prp24’s absence. Post-transcriptional processing facilitates Lsm2-8 association while the presence of Prp24 promotes both recruitment and retention of the complex. This suggests that efficient U6 snRNP assembly could depend on kinetic selection of Lsm2-8 binding to 3’-end modified or Prp24 bound U6 snRNAs in order to discriminate against association with other RNAs.

## INTRODUCTION

In eukaryotes, the removal of introns from pre-mRNA is accomplished by a dynamic molecular machine known as the spliceosome. This machine is partly composed of five small nuclear ribonucleoproteins (snRNPs) named U1, U2, U4, U5, and U6, which each consist of small nuclear RNA (snRNA) molecules and associated protein factors. Throughout the splicing process, the spliceosome undergoes a precisely orchestrated series of assembly, activation, catalysis, dissociation, and recycling steps involving the snRNP particles (1–3). Among the snRNAs, U6 is the most highly conserved and the most dynamic (4, 5). For splicing to occur, U6 snRNPs containing the U6 snRNA, Prp24 protein, and Lsm2-8 protein heteroheptamer must first associate with the U4 snRNP and form base-pairing interactions between the U4 and U6 snRNAs (5). The U4/U6 di-snRNP must then associate with the U5 snRNP to form the U4/U6.U5 tri-snRNP, and Prp24 must dissociate at some point during this process (6). The tri-snRNP is subsequently integrated into the spliceosome and then drastically remodeled: the U4 snRNP and Lsm2-8 complexes are released to allow base pairing between the U2 and U6 snRNAs, between the U6 snRNA and the intron, and intramolecular base pairing within U6 itself. When splicing concludes, U6 snRNA must be released from the spliceosome by translocation of the Prp43 helicase along the RNA in the 3’→5’ direction (7–9). At some point, U6 snRNA then re-associates with Prp24 and Lsm2-8 to regenerate the U6 snRNP.

Initial biogenesis of the U6 snRNP follows a similar pathway involving Prp24 and Lsm2-8 (5). However, the nascent U6 snRNA must also be post-transcriptionally modified. U6 snRNA is generated by RNA polymerase (RNAP) III, which terminates transcription in response to a poly- U tract found at the 3’ end of U6 (10). After transcription, the 3’ terminal diol of U6 is initially bound by the protein chaperone Lhp1, providing protection against degradation (11). Following the association of Prp24, Lhp1 is released, and the 3’ end is exposed for processing by the Usb1 exonuclease (12). Usb1 removes a terminal uridine from U6 while generating a 3’ phosphate at the new terminus (12, 13). While not strictly essential for Lsm2-8 binding, this processing increases the equilibrium affinity of Lsm2-8 for U6 by 3- to 7-fold (12).

The assembly of the *Saccharomyces cerevisiae* (*S. cerevisiae*) U6 snRNP has been reconstituted from purified components *in vitro* and is highly efficient (12, 14, 15). This represents a powerful system for dissecting how RNA and proteins dynamically interact with one another during molecular assembly and for connecting *in vivo* genetic results to quantifiable changes in structure, kinetic, or thermodynamic stabilities. While several *in vitro* studies have elucidated the equilibrium parameters that describe binding between the U6 snRNA and its associated proteins (*i.e., K*_D_’s) (12, 14, 16–18), there is little information about the kinetic details of U6 snRNP assembly starting either from a nascent transcript or the 3’-end modified snRNA. In cells, it is possible that kinetic pathways dominate how U6 snRNPs are assembled since competing, non-reversible pathways of RNA export, degradation, and assembly into larger splicing complexes may prevent equilibrium conditions from being established. Indeed, it is well-known that many steps in gene expression do not occur at equilibrium in cells (19).

In this study, we have analyzed the kinetic properties of yeast U6 snRNP assembly *in vitro* using purified components and colocalization single molecule spectroscopy (CoSMoS). Single molecule experiments like CoSMoS offer distinct advantages compared to ensemble assays by enabling the detection of transient intermediates, facilitating the identification of alternative assembly pathways, and visualizing complexes in real-time as they form rather than with discontinuous experiments such as electrophoretic mobility shift assays (EMSAs) (20, 21). By studying the kinetics of Lsm2-8 complex binding with U6 snRNA as a function of RNA 3’-end composition and presence or absence of Prp24, we show that post-transcriptional processing of U6 significantly changes Lsm2-8 binding kinetics. Addition of Prp24 appears to optimize this interaction, further stabilizing the complex. In particular, the SNFFL-box motif in the C-terminal domain of Prp24 can modulate U6’s accessibility for Lsm2-8 binding depending on the RNA’s post-transcriptional modification status. Together our results suggest that 3’-end modification of U6 may increase the efficiency of U6 snRNP assembly by providing access to an alternate pathway in which Lsm2-8 can rapidly bind prior to Prp24 association.

## RESULTS

### Fluorophore-labelled Lsm2-8 binds 3’-end modified RNAs

For imaging single-molecules of yeast Lsm2-8, we constructed an Lsm2-8 expression plasmid in which Lsm8 is genetically fused at the C terminus to the fast SNAP tag (Lsm2-8^fSNAP^; **Fig.1A**, **B**). We have previously shown that a similar C-terminal fSNAP fusion to Lsm8 is functional *in vivo* and assembles into spliceosomes (22). We expressed and purified assembled Lsm2-8^fSNAP^ complexes from *E. coli*. Lsm2-8^fSNAP^ was then labelled SNAP-Surface 549 (DY549), and the excess dye was removed by gel filtration chromatography (**Fig. S1A**). UV-vis spectroscopy was used to determine the extent of fluorophore labelling (∼80%).

**Figure 1.**
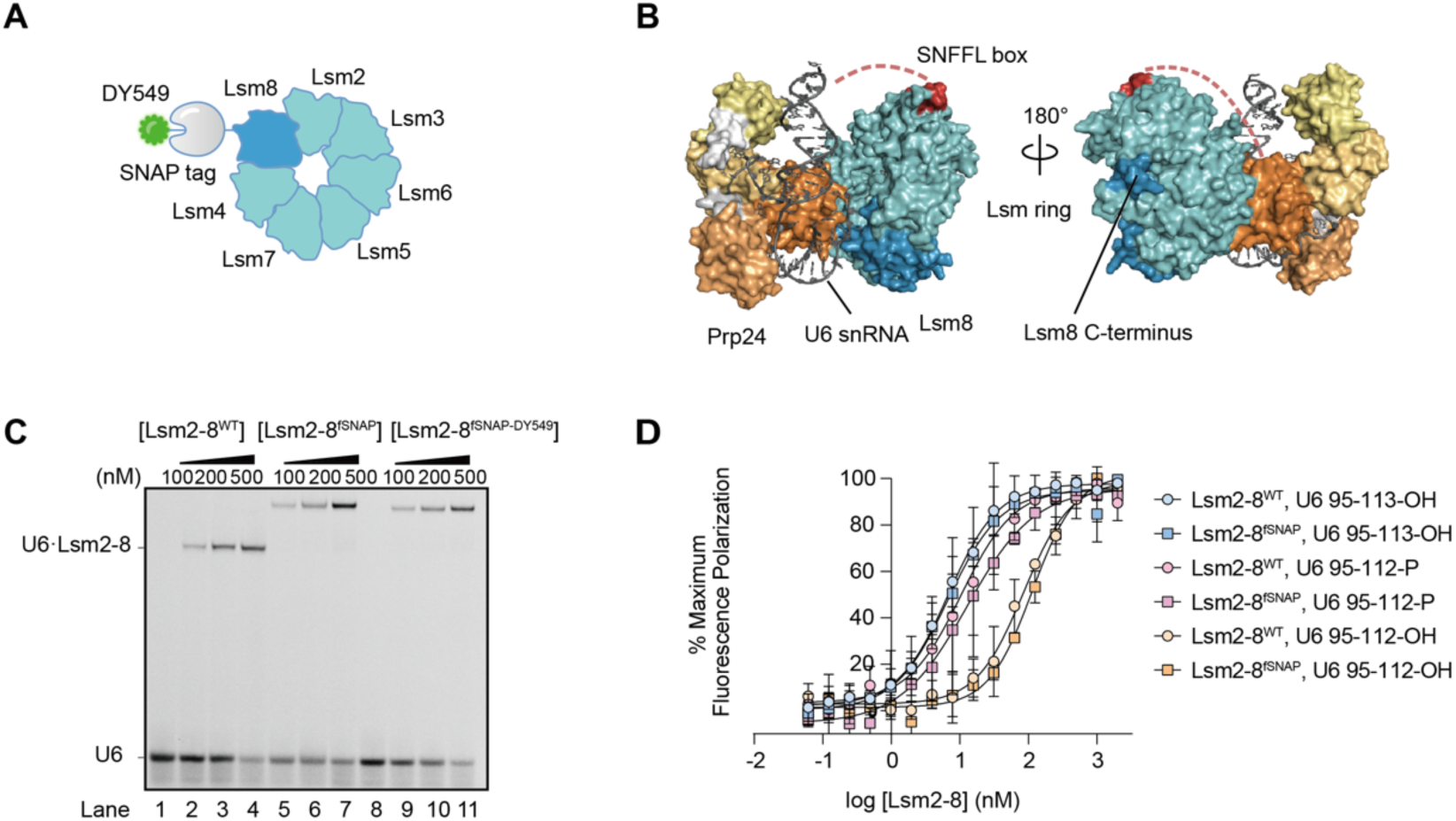
Fluorophore-labeled yeast Lsm2-8 binds 3’ end modified RNAs. **(A)** Cartoon structure of Lsm2-8^fSNAP_DY549^. The SNAP tag is attached to the C-terminus of Lsm8 via a flexible linker. **(B)** Crystal structure of the yeast U6 snRNP highlighting the position of Lsm8 (blue), Lsm8 C-terminus (purple blue), and the Prp24 C-terminal SNFFL box (red) (PDB ID: 5VSU). **(C)** EMSA analysis of Lsm2-8^fSNAP^ and Lsm2-8^fSNAP_DY549^ binding ability relative to WT protein to U6 RNAs. **(D)** Fluorescence polarization binding data comparing Lsm2-8^WT^ and Lsm2-8^fSNAP^ binding to U6 RNA fragments (nt 95-112 or -113) with wither diol or 3’ phosphate end modifications.

We tested the impact of fSNAP labelling on Lsm2-8 binding to U6 RNAs. EMSA assays confirmed that the fluorophore-labelled Lsm2-8^fSNAP^ bound an RNA oligo representing U6 nucleotides (nt) 95-113 with a terminal 2’, 3’ diol (U6 95-113-OH) similarly to the untagged complex (**Fig. 1C**). We then used fluorescence polarization assays to study the affinities of unlabeled Lsm2-8^fSNAP^ complexes for RNAs with different modification states of U6 (**Fig. 1D**). Like Lsm2- 8^WT^, Lsm2-8^fSNAP^ showed the highest affinities for RNAs with either a five uridine 3’ end (U6 95- 113-OH) or with a four uridine 3’ end containing a 3’ phosphate (U6 95-112-P; **Supplementary Table 1**). RNAs containing four uridines and a 2’, 3’ diol at the 3’ end (U6 95-112-OH) bound both complexes with lower affinity. From these assays we conclude that the labelled Lsm2-8^fSNAP^ complex interacts similarly with U6 RNA as does the untagged version.

### Single Molecule Assays of U6 RNA Binding Recapitulate Ensemble Observations

To image Lsm2-8^fSNAP^ proteins associated with U6 snRNAs, we used 2-color CoSMoS assays (23). In these experiments, full-length, Cy5-labeled U6 RNAs (nts 1-113; 1-113-OH) were prepared by splinted ligation (**Fig. S1B**) and then immobilized to passivated glass slides with biotin-streptavidin. Interactions with DY549-labeled Lsm2-8^fSNAP^ could then be visualized by observing colocalization between the red (632 nm) laser- and green (532 nm) laser-excitable fluorophores on the RNA and Lsm2-8^fSNAP^, respectively (**Fig. 2A**). Data were collected using a micromirror multi-wavelength total internal reflection fluorescence (TIRF) microscope (24, 25).

**Figure 2.**
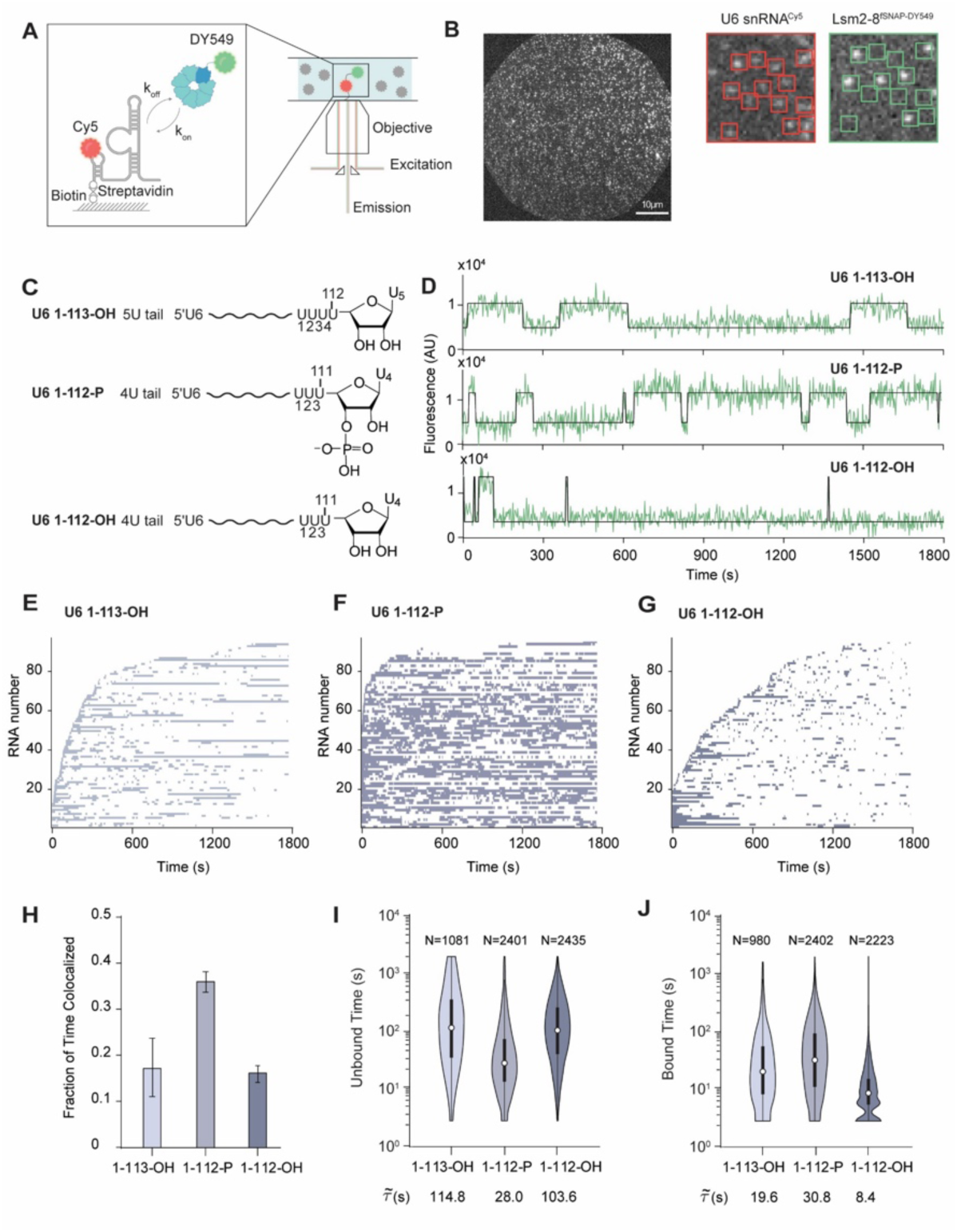
U6 RNA 3’ end modification impacts Lsm2-8^fsnap^ binding kinetics. **(A)** Schematic representation of the CoSMoS assay for monitoring Lsm2-8^fSNAP^ ^DY549^ binding to immobilized, Cy5-labeled U6 RNAs. **(B)** Left, representative micrographs showing individual U6 snRNA molecules tethered to the slide surface in a field of view (FOV). Right, zoom-in of a portion of the FOV showing individual U6 RNA molecules (red boxes on the left) and colocalized Lsm2-8^fSNAP^ molecules (bright spots inside the green boxes on the right). **(C)** Chemical structures of the RNAs used in the CoSMoS assays highlighting the structure of the RNA 3’ ends. **(D)** Representative fluorescence trajectories showing Lsm2-8^fSNAP^ ^DY549^ binding events to the RNAs. **(E-G)** Rastergrams illustrating the binding of Lsm2-8 to 90 different molecules of U6 1-113-OH (E), U6 1-112-P (F), and U6 1-112-OH (G), sorted based on the time of first Lsm2-8^fSNAP^ ^DY549^ binding event detection. **(H)** Average fraction of time U6 molecules spend colocalized with Lsm2-8^fSNAP^ ^DY549^. Bar heights represent the average from three experimental replicates and bars represent ±SD. **(I, J)** Violin plots representing the dwell time distributions for unbound (I) and bound (J) states across three different RNAs. Each violin plot is overlaid with a box plot that indicates the median (horizontal line), interquartile range (IQR, box), and whiskers extending to 1.5 times the IQR. The numbers above each violin plot denote the number of lifetimes included in the respective distribution. The median dwell times are shown below each plot.

In a specific field of view, we simultaneously tracked binding events occurring on hundreds of U6 1-113-OH RNA molecules (**Fig. 2B**). We frequently observed spots of fluorescence from Lsm2-8^fSNAP^ dynamically colocalizing with tethered RNA locations—indicating that Lsm2-8 association with the RNAs is reversible. As a control, if RNA was omitted from the surface, we observed significantly fewer (∼15-fold lower) Lsm2-8^fSNAP^ binding events (**Fig. S2**). This suggests that colocalization events observed between Lsm2-8^fSNAP^ and the immobilized RNAs represent specific RNA/protein interactions. To test if these interactions required the free 3’ end of the U6 RNA, we first incubated the immobilized RNAs with the yeast Lhp1 protein, which tightly binds U6 RNAs containing a terminal 3’ diol. We predicted that this should sterically block the association of the fluorescent Lsm2-8^fSNAP^ complex with U6 1-113-OH. This prediction was correct and addition of Lhp1 reduced the number of colocalized Lsm2-8^fSNAP^ binding events to background levels (**Fig. S2**). Finally, we used yeast Prp24 protein to release Lhp1 from the immobilized U6 RNAs and restore Lsm2-8^fSNAP^ binding (**Fig. S2**). Together, these results indicate that Lsm2-8^fSNAP^ molecules colocalize with immobilized U6 RNAs depending on the accessibility of the RNA 3’ end and that Prp24 can release Lhp1 from immobilized U6 RNAs. We conclude that these single molecule assays recapitulate key features of the U6 snRNP assembly pathway defined by bulk experiments (12).

### 3’ End Modification of U6 Changes Lsm2-8 Occupancy and Interaction Kinetics

In yeast, U6 snRNAs are synthesized by RNAP III, and transcription termination generates heterogeneously sized U6 tails ranging from 4-8 nt (5). Processing by Usb1 leads to the reduction of the U-tail length by one nucleotide and leaves behind a terminal 3’ phosphate group (12). In order to investigate the influence of different tail lengths and chemical groups on the binding behavior of Lsm2-8^fSNAP^, we carried out CoSMoS assays with Lsm2-8^fSNAP^ on three different types of full length, immobilized U6 RNAs: a mimic of the unprocessed form (U6 1-113-OH), a mimic of the product of yUsb1 processing (U6 1-112-P), and a final form lacking both one uridine in the 3’ tail and the 3’ phosphate group (U6 1-112-OH) (**Fig. 2C**).

Representative fluorescence trajectories of Lsm2-8^fSNAP^ fluorescence colocalized with individual U6 RNAs are shown in **Fig. 2D**. These trajectories suggest that the modification status of the U6 RNA 3’ end can change Lsm2-8^fSNAP^ interaction dynamics: fewer and shorter events are observed with the U6 1-112-OH RNA relative to the other two. To provide overviews of binding events occurring on many different RNAs, we created rastergrams where the colocalization of Lsm2-8^fSNAP^ is represented by a purple segment and its absence by a white segment for 90 different RNA molecules. These rastergrams were further sorted by the timing of the first Lsm2- 8^fSNAP^ binding event, with RNAs binding Lsm2-8^fSNAP^ earlier being assigned a lower number on the y-axis (**Fig. 2E-G**).

From the rastergrams, it is evident that many more binding events are observed on the U6 1-112-P RNA relative to the others. In contrast, binding events to an RNA of the same length but lacking the 3’ phosphoryl modification (U6 1-112-OH) are both shorter and less frequent. Increasing the RNA tail length restores some long-lived binding events (U6 1-113 OH) but not the event density seen with the phosphoryl modification. Together, the rastergrams indicate that the U6 RNA mimicking modification by Usb1 has increased interaction with Lsm2-8^fSNAP^.

To quantify these changes more accurately, we first determined the total fractions of experimental times each RNA was colocalized with Lsm2-8^fSNAP^ (**Fig. 2H**). As expected from the rastergrams, U6 1-112-P RNAs spent nearly twice as often colocalized with Lsm2-8^fSNAP^ proteins as did the other RNAs.

We then analyzed the distributions of dwell times for binding events as well as the times in between binding events (**Fig. 2I, J**). These distributions show that Lsm2-8 complexes tended to bind the U6 1-112-P RNA more quickly (lower unbound times; **Fig. 2I**) than either the 1-113- OH or 1-112-OH RNAs. Once bound, Lsm2-8 complexes tended to remain associated with the 1- 113-OH and 1-112-P RNAs longer relative to the 1-112-OH RNA. To analyze the dwell times in more detail, we fit each distribution of dwell times to an equation containing two exponential terms to yield characteristic lifetimes for the bound (1_bound_) and unbound (1_unbound_) states (**Supplementary Tables 2**, **3**). For the unbound times, we observed a multi-exponential distribution for each RNA with characteristic short- and long-1_unbound_ parameters. In comparing the unmodified and modified U6 RNA mimics (U6 1-113-OH and U6 1-112-P, respectively), the most significant differences are found in a large increase in the amplitude of the short-1_unbound_ parameter (increasing from 0.50 to 0.81) and a decrease in the value of 1_unbound_ from ∼86 to 32 s due to the modification. One interpretation of these data is that U6 end modification increases both the rate of Lsm2-8 binding (as observed in the decrease 1_unbound_) and the likelihood of the RNA being able to bind Lsm2-8 rapidly (as observed by the increase in amplitude).

We also analyzed the distributions of bound dwell times (**Supplementary Table 3**). In all cases, we observed a long-lived component of ∼200-300 s. To determine if the lifetimes of the longest-lived events could be shortened due to photobleaching, we analyzed the photobleaching rate of an immobilized fSNAP protein labeled with DY549 under the same conditions (**Fig. S3**). For the photobleaching control, the labeled SNAP protein had a characteristic lifetime of ∼2682 s, at least 10-fold longer than the lifetimes of the Lsm2-8^fSNAP^ proteins when associated with U6. Therefore, we assume that photobleaching had a minimal impact on the observed lifetimes of the longest-lived Lsm2-8^fSNAP^ binding events.

In contrast with the analysis of unbound dwell times, the characteristic 1_bound_ parameters for the U6 1-113-OH and U6 1-112-P RNAs were quite similar to one another. In both cases, the fits were best described with two exponential terms. The amplitudes of the short- and long-lived parameters were only modestly higher (0.20 vs. 0.31) for the long-lived parameter for U6 1-112-P. This suggests that while 3’ end modification can increase Lsm2-8 occupancy on U6, the origin of this effect stems mostly from facilitating protein association rather than just stabilization of the bound state. Interestingly, the U6 1-112-OH RNA showed a dramatic decrease in bound state lifetimes with an amplitude of the long-lived parameter of only 0.01. The vast majority of binding events observed on that RNA were short-lived with a characteristic 1_bound_ of ∼12 s. This RNA also showed an increase in the characteristic times for 1_unbound_ (**Supplementary Table 2**). Lack of Lsm2-8^fSNAP^ occupancy on the U6 1-112-OH RNA originates both from difficulties in recruiting the protein complex to the RNA and stabilization of the complex once bound.

### Kinetic Mechanism for Lsm2-8 Recruitment to Modified U6 RNAs

We further analyzed the single molecule data for Lsm2-8^fSNAP^ binding the U6 1-112-P RNA to determine a kinetic mechanism that fit the observed dwell times. We carried out this analysis only with the U6 1-112-P RNA since it likely represents the predominant species for Lsm2-8 recruitment during U6 snRNP recycling from spliceosomes. We collected single molecule data at various concentrations of Lsm2-8^fSNAP^ (1, 3, and 10 nM), and then globally fit the data to various kinetic schemes containing 2, 3, or 4 states using QuB software. We determined the best fit to be to a four-state model (**Fig. S4**). Lsm2-8 binding predominantly occurs in a single step with a *k_on_* of 1.94 × 10^7^ M^-1^·s^-1^ and *k_off_* = 0.045 s^-1^. However, either or both U6 and Lsm2-8 can rarely transition to a state from which the U6/Lsm2-8 complex cannot form directly. Additionally, the U6/Lsm2-8 complex itself can rarely transition to a state from which Lsm2-8 cannot directly dissociate. It is possible that these rare states represent alternate conformations of the protein or RNA that preclude Lsm2-8 association or dissociation.

### Prp24 Facilitates Lsm2-8 Recruitment on Unmodified RNAs

Since binding of Prp24 to displace Lhp1 is a pre-requisite for subsequent Lsm2-8 binding on unmodified U6 RNAs (12, 26, 27), we next investigated the influence of Prp24 on Lsm2-8^fSNAP^ interactions with U6 (**Fig. 3A**). In these experiments, we added a solution containing both Prp24 and Lsm2-8^fSNAP^ to slides containing immobilized U6 molecules and monitored Lsm2-8^fSNAP^ binding to the RNAs. One caveat of these experiments is that we did not monitor Prp24 binding directly; however, it was added at a concentration of 100 nM, ∼5-fold higher than the reported *K*_D_ 21nM (14).

**Figure 3.**
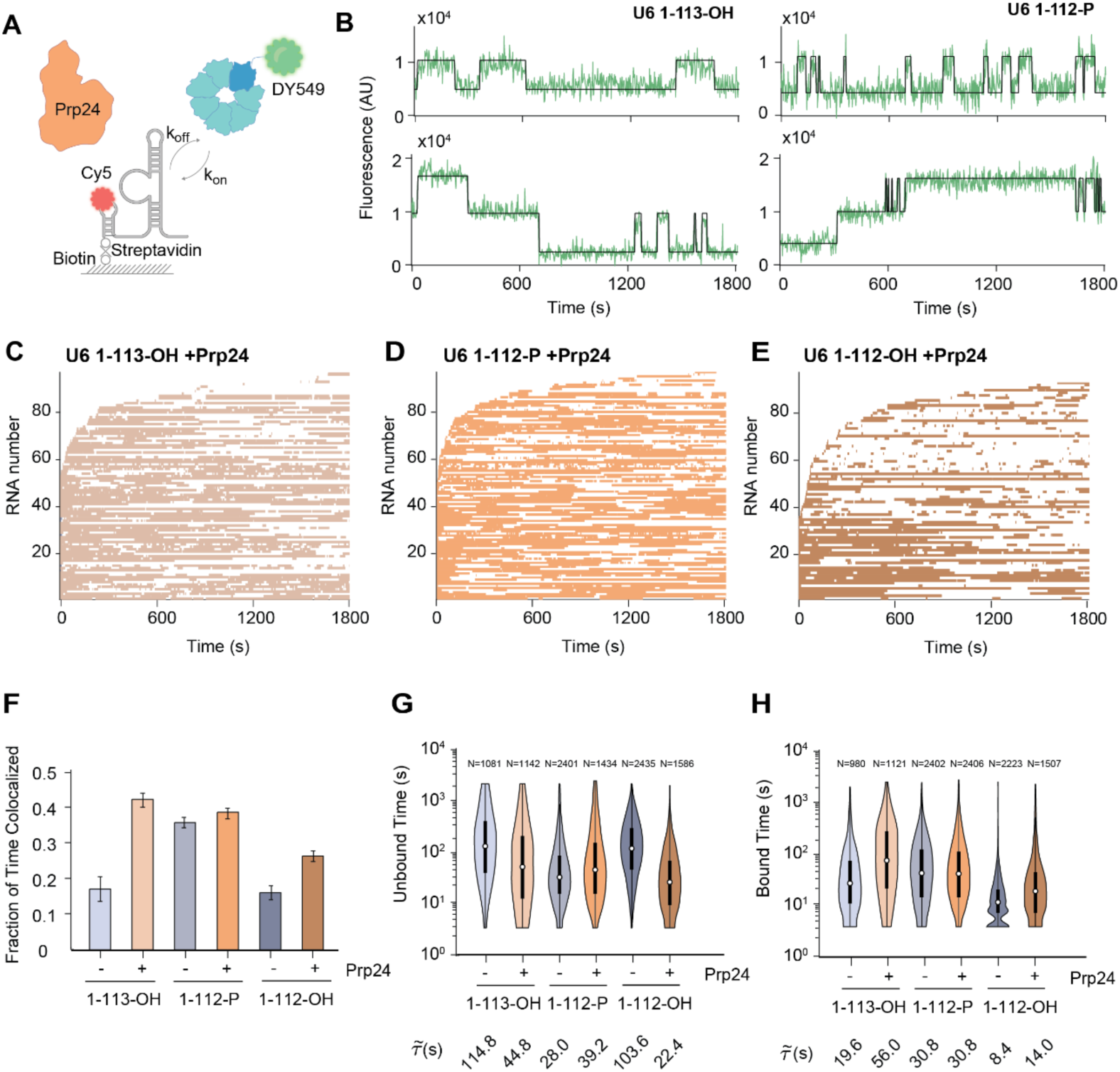
Prp24 enhances Lsm2-8^fSNAP^ binding kinetics to unmodified RNAs. **(A)** Schematic representation of the CoSMoS assay for monitoring Lsm2-8^fSNAP^ ^DY549^ binding in the presence of unlabeled Prp24. **(B)** Representative fluorescence trajectories showing single Lsm2-8^fSNAP^ ^DY549^ (top traces) or double (bottom traces) to RNAs in the presence of Prp24. **(C-E)** Rastergrams illustrating the binding of Lsm2-8 to 90 different molecules of U6 1-113-OH (C), U6 1-112-P (D), and U6 1-112-OH (E), sorted based on the time of first Lsm2-8^fSNAP^ ^DY549^ binding event detection. **(F)** Average fraction of time U6 molecules spend colocalized with Lsm2-8^fSNAP^ ^DY549^ in the presence of Prp24. Bar heights represent the average from three experimental replicates and bars represent ±SD. Data for the absence of Prp24 is replicated from Fig. 2H to facilitate comparison. **(G, H)** Violin plots representing the dwell time distributions for unbound (G) and bound (H) states across three different RNAs. Each violin plot is overlaid with a box plot that indicates the median (horizontal line), interquartile range (IQR, box), and whiskers extending to 1.5 times the IQR. The numbers above each violin plot denote the number of lifetimes included in the respective distribution. The median dwell times are shown below each plot. Data for the absence of Prp24 is replicated from Fig. 2I, J to facilitate comparison. For data shown in panels C-H, only RNAs containing single binding events were analyzed.

When we analyzed individual fluorescence trajectories, we noticed that while the majority RNAs bound only one Lsm2-8^fSNAP^ complex at a time, a small subset showed stepwise changes in fluorescence consistent with the presence of at least two Lsm2-8^fSNAP^ complexes associated with the same RNA (**Fig. 3B**). To simplify our analysis, we first analyzed only the subset of RNAs that bound a single Lsm2-8 ^fSNAP^ complex at a time. A rastergram analysis of these RNAs revealed that addition of Prp24 increases the association of Lsm2-8^fSNAP^ with U6 regardless of the 3’ end modification (**Fig. 3C-3E**). Lsm2-8 exhibited similar binding patterns on both U6 1-113-OH and 1- 112-P, while binding on 1-112-OH appeared weaker but still stronger than was observed in the absence of Prp24 (**Fig. 2F**).

When we quantified the extent of Lsm2-8^fSNAP^ occupancy on RNAs in the presence of Prp24, we saw little change in occupancy with U6 1-112-P (**Fig. 3F**). However, Prp24 increased the occupancy fractions for both the 1-113-OH and 1-112-OH RNAs to levels more closely resembling that observed for 1-112-P. When comparing the distributions of dwell times (**Fig. 3G, H**), it is apparent that Prp24 can perturb both unbound and bound times for the 1-113-OH and 1- 112-OH RNAs. In contrast, the distributions of times observed on the 1-112-P RNA were similar regardless of the presence or absence of Prp24.

When comparing the characteristic 1_unbound_ and 1_bound_ times for the unmodified (1-113-OH) and modified (1-112-P) RNA mimics in the presence of Prp24, the effect of Prp24 is most pronounced in a change in the long-lived 1_bound_ parameter for U6 1-113-OH (**Supplementary Table 3**). The amplitude of this parameter doubles from ∼0.20 to 0.55 due to the presence of Prp24, and the associated characteristic lifetime increases from ∼183 to 290 s. Similarly, parameters for the unbound state for U6 1-113-OH show a reduction in the times between binding events (**Supplementary Table 2**). Overall, the parameters for the Lsm2-8^fSNAP^ binding to U6 1- 113-OH in the presence of Prp24 more closely resemble those for binding to U6 1-112-P than when Prp24 is absent. The U6 1-112-OH RNA is also impacted by Prp24 with increased binding due to changes in both the amplitude of the long-lived bound state (∼10-fold increase) and decreased times between binding events. In summary, Prp24 can facilitate the recruitment of Lsm2-8 to unmodified RNAs and the longest-lived complexes between Lsm2-8 and U6 have similar lifetimes independent of the 3’ end modification when Prp24 is present (300-400 s).

### The SNFFL Box is Required for Prp24 to Promote Lsm2-8 Binding to Unmodified U6 RNAs

Previous studies have shown that the SNFFL box of Prp24 interacts with Lsm2-8 (**Fig. 1B**). We wondered if a U6-Prp24 complex could present multiple surfaces for Lsm2-8 interaction, one of which could be dependent on the SNFFL box(15). These multiple surfaces could result in the double binding events we observed on some RNAs in the presence of Prp24 (**Fig. 3B**). To test this, we purified a Prp24 mutant lacking the SNFFL box. In comparison with WT Prp24, Prp24^ΔSNFFL^ is less efficient at forming U6-Prp24 and U6-Prp24-Lsm2-8 complexes by EMSA (**Fig. 4A**).

**Figure 4.**
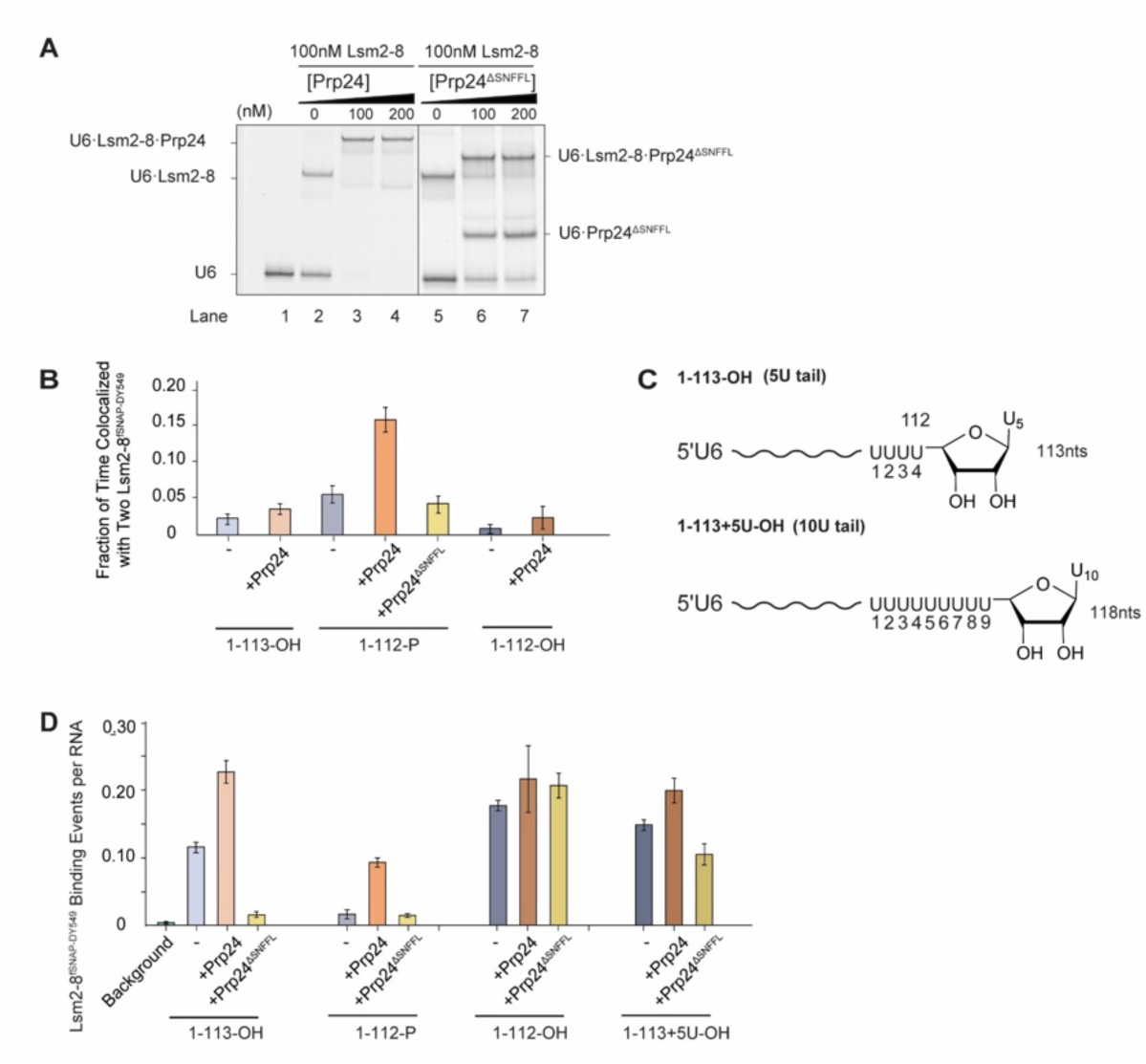
The Prp24 SNFFL box is required for recruitment of Lsm2-8^fSNAP^ to unmodified U6 RNAs. **(A)** EMSA analysis of Lsm2-8/Prp24/U6 complex formation using EMSA using Prp24 containing (lanes 3-4) or lacking (lanes 6-7) the SNFFL-box domain. The concentrations of Cy5-labeled U6 oligos and Lsm2-8 were maintained at a constant 2nM and 100nM, respectively. **(B)** Average fractions of time U6 RNAs are simultaneously bound by two Lsm2-8^fSNAP^ ^DY549^ complexes. **(C)** Chemical structures of the U6 1-113-OH and 1-113+5U-OH RNAs, highlighting the RNA 3’ ends. **(D)** Relative numbers of Lsm2-8^fSNAP^ ^DY549^ binding events per U6 RNA in the presence or absence of Prp24 and Prp24^ΔSNFFL^. In panels B and D, bar heights represent the average from three experimental replicates and bars represent ±SD.

We then determined if use of the Prp24^ΔSNFFL^ protein would change the frequencies at which we observed double binding events of Lsm2-8^fSNAP^ on U6 RNAs in the CoSMoS assay. We tested this with the U6 1-112-P RNA since double binding events were much more rarely observed with the other RNAs (**Fig. 4B**). With U6 1-112-P, RNAs spent nearly 15% of the total experimental time associated with more than one Lsm2-8^fSNAP^ protein in the presence of Prp24 compared to ∼5% in its absence. However, when Prp24^ΔSNFFL^ was used, this increase was not observed and observations of double occupancy in the presence of Prp24^ΔSNFFL^ were similar in frequency as to when Prp24 was omitted altogether. These data suggest that more than one Lsm2-8 complex can be recruited to a single U6 RNA at one time via the SNFFL domain of Prp24.

Finally, we tested if the SNFFL domain would have any impact on Prp24’s ability to recruit Lsm2-8^fSNAP^ to unmodified U6 RNAs 1-112-OH or 1-113-OH. Much to our surprise, the addition of Prp24^ΔSNFFL^ prevented Lsm2-8^fSNAP^ binding to the U6 1-112-OH and 1-113-OH RNAs but not to U6 1-112-P (**Fig 4C**). This was unexpected since we observed strong binding of Lsm2-8^fSNAP^ to both RNAs in the presence of full length Prp24. In fact, Lsm2-8^fSNAP^ binding to the 1-113-OH RNA in the presence of Prp24^ΔSNFFL^ was less than if Prp24 was omitted entirely. This suggests Prp24^ΔSNFFL^ can inhibit Lsm2-8 from binding cis-diol terminated U6 RNAs.

A possible mechanism to account for this could be sequestration of the 3’ end of U6 by Prp24^ΔSNFFL^ when it contains a cis-diol but not a terminal phosphate. This sequestration then prevents Lsm2-8 binding. To test this hypothesis, we constructed a cis-diol terminated U6 RNA with an extended 3’ end (1-113+5U-OH) in an attempt to attenuate the hindrance by Prp24^ΔSNFFL^ (**Fig. 4D**). In agreement of our hypothesis, Lsm2-8 exhibited increased binding to this longer RNA suggesting that the RNA 3’ end is now accessible (**Fig. 4D**). Together, our findings indicate that not only can the SNFFL box function to recruit Lsm2-8 complexes to U6 but that it is also essential for permitting Lsm2-8 recruitment to unmodified RNAs bound by Prp24.

## DISCUSSION

For efficient pre-mRNA splicing, it is important that the U6 snRNA be rapidly recycled from spliceosomes into U6 snRNPs and that the snRNA be protected from degradation (5). Previous work established a pathway by which yeast U6 snRNP can be post-transcriptionally modified on its 3’ end and assembled *in vitro* (12). In this work, we show that 3’ end modification can alter the kinetic pathways of assembly. In agreement with ensemble data, we observed a much higher degree of binding of Lsm2-8 to single U6 RNAs that mimic the processed form (U6 1-112-P). The origin of this effect likely stems from the presence of the 3’ phosphate since a shorter RNA containing a diol (U6 1-112-OH) did not stimulate binding. Dwell time analysis indicates that the increase in binding is primarily due to the RNA modification facilitating Lsm2-8 association (shorter 1_unbound_ times) rather than just stabilization of the bound state. Prp24 can moderate the impact of 3’ end modification and allow for efficient recruitment of Lsm2-8 on both processed and unprocessed RNAs. The increase in Lsm2-8 occupancy due to Prp24 can arise from both facilitating association and slowing dissociation. Interestingly, we observe that some U6 RNAs can recruit two Lsm2-8 complexes simultaneously in the presence of Prp24 and that this is dependent on the Prp24 SNFFL-box domain. This domain is also required for Prp24 to recruit Lsm2-8 to unmodified RNAs since Prp24 molecules lacking the SNFFL-box can actually inhibit Lsm2-8 binding. We do not know if Lsm2-8 complexes bound to SNFFL-box but not yet associated with U6 can be directly transferred to the RNA 3’ end.

Together our results indicate that post-transcriptional modification of U6 facilitates snRNP assembly by opening an alternate route of Lsm2-8 acquisition (**Fig. 5**). In the absence of modification, Lsm2-8 binding would be highly dependent on the presence of Prp24. For this pathway, we assume that Prp24 binds U6 first and subsequently recruits Lsm2-8. However, we did not simultaneously monitor Prp24 and Lsm2-8 recruitment, and it is possible that a pre-formed Prp24/Lsm2-8 complex may also be involved in snRNP assembly. U6 snRNA modification opens a second pathway in which efficient and stable Lsm2-8 binding occurs in the absence of Prp24.

**Figure 5.**
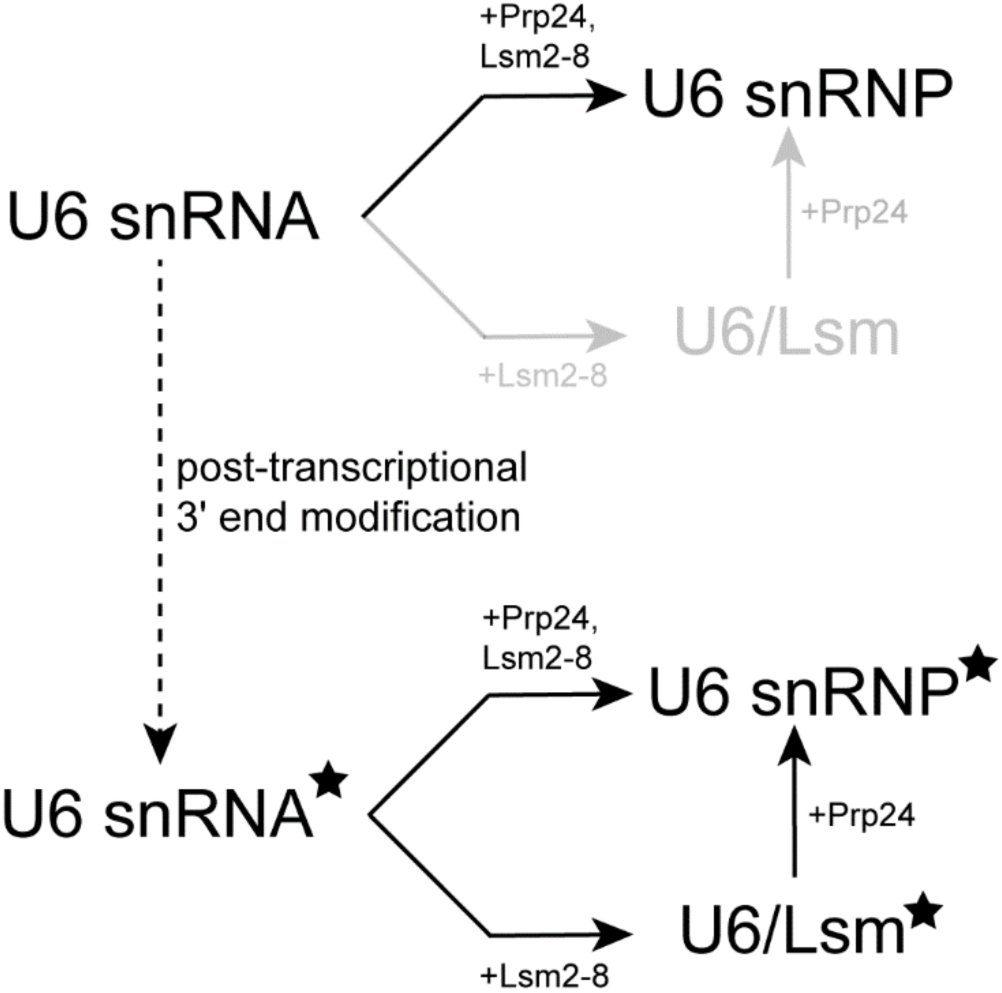
U6 RNA 3’ end modifications permits an alternate assembly route for the U6 snRNP. The unmodified U6 snRNA depends on interaction with Prp24 for Lsm2-8 recruitment while U6 modification (3’ phosphate modification is denoted by the black star) permits Prp24-dependent and -independent Lsm2-8 recruitment.

Both Prp24-dependent and independent pathways may be functionally relevant for splicing *in vivo*. In the case of *de novo* U6 snRNP assembly from an unprocessed transcript, Prp24 dependence likely assures that the Lsm2-8 complex assembles specifically on U6 snRNAs rather than on any transcript harboring a 3’ polyU sequence. Notably, such transcripts are abundant in the cell since all RNAPIII transcripts terminate with polyU including tRNAs and 5S rRNAs. In terms of recycling U6 snRNA from spliceosomes at the conclusion of splicing, direct recruitment of Lsm2-8 to modified U6 snRNA may facilitate rapid re-assembly of the snRNP while the spliceosome is being dismantled and before Prp24 has the opportunity to bind. In addition to making re-assembly more efficient, this may also be important for protecting U6 snRNA from nucleases by ensuring fast sequestration of the RNA 3’ end (26, 28).

It is interesting to note that the related Lsm1-7 complex from *S. pombe* can tightly bind polyU tracts on RNAs lacking modifications and in the absence of protein cofactors (Pat1) (29). Even though these complexes share 6 out of 7 subunits, their functions are very different: Lsm1- 7 association marks mRNAs for degradation but Lsm2-8 association is needed for U6 snRNP assembly. While we have not carried out a detailed kinetic analysis of Lsm1-7/RNA interactions, this observation suggests that it may be equally important for the cell to prevent Lsm2-8 (or Lsm1- 7) complex assembly on the wrong RNAs as it is to facilitate assembly on U6. Dependence of the Lsm2-8 binding kinetics on both 3’ end modification and the presence of Prp24 could provide two independent layers of proofreading via kinetic selection to facilitate preferential recruitment to the U6 snRNA.

## EXPERIMENTAL PROCEDURES

### Cloning and Mutagenesis

The fSNAP variant of yeast Lsm2-8 was created by first removing the Lsm8 sequence from the pQLink-Lsm2-8 plasmid (a gift from Yigong Shi; (30)) by restriction enzyme digestion with SacI and NheI. A double-stranded DNA fragment coding for a c-terminal fusion of the fSNAP tag to Lsm8 was synthesized (ThermoFisher, GeneArt Synthetic Gene) and ligated into those same restriction sites. To ensure the accuracy of recombinant plasmids and deletion mutants, DNA sequencing was performed for the whole plasmid.

### Protein Expression and Purification

Prp24 was purified as described previously (12). Briefly, proteins were expressed in E. coli BL21 (DE3) pLysS cells in LB medium after induction by addition of isopropyl β-D-1- thiogalactopyranoside (IPTG) to a final concentration of 1 mM for 20h at 16°C. Proteins were then purified by Ni-NTA agarose chromatography followed by removal of the his-tag with TEV protease and cation-exchange chromatography as described except that a HiTrap SP FF (GE Healthcare) column was used.

Lhp1 was also purified as previously described (12). Lhp1 protein was expressed in E. coli BL21 (DE3) pLysS cells in LB medium after induction by addition of IPTG (1 mM) for 20h at 16°C. Lhp1 was then purified by Ni-NTA agarose chromatography followed by removal of the histag with TEV protease and cation-exchange chromatography as described except that a HiTrap S (GE Healthcare) column was used.

Lsm2-8 and Lsm2-8^fSNAP^ were purified using a previously described protocol for Lsm2-8 (12). Lsm proteins were expressed in E. coli BL21 (DE3) pLysS cells in LB medium after induction by addition of IPTG (1 mM) for 20h at 37°C. Lsm protein complexes were then purified by Ni-NTA agarose and GST-affinity chromatography followed by removal of the histag with TEV protease and anion-exchange chromatography as described.

For all proteins, the final concentrations were determined by using calculated extinction coefficients (ProtParam, (31)) and measuring the absorbance at 280 nm.

### Fluorophore Labeling of Lsm2-8^fSNAP^

SNAP-Surface® 549 dye (S9112S, New England BioLabs) in DMSO was added to a protein solution (500 nM; 500 µL) at a dye ratio 1.25 dye:1 Lsm2-8^fSNAP^ complex in SEC buffer (25 mM HEPES-KOH pH 7.9, 50 mM KCl, 1 mM TCEP, 10% v/v glycerol). The reaction tube was then incubated in darkness at room temperature (RT) for 1 h. Excess dye was removed by gel filtration chromatography using G-25 Sephadex (Sigma) equilibrated in SEC buffer. For the labeled proteins, the labeling efficiency was determined by the concentrations of both dye and protein in the final product measured by the absorbance at 280 nm and 550nm.

### RNA Preparation

RNA oligos were purchased from Integrated DNA Technologies (IDT). Sequences and modifications are described in **Supplemental Table S4.**

Full-length U6 RNAs (112 or 113 nt in length) containing a 3’ *cis-diol* were prepared as previously described by splinted RNA ligation using T4 RNA ligase 2 (12). For CoSMoS assays, the 5’ piece of the U6 RNA encoding nt 1-12 also contained a 5’ biotin and internal Cy5 fluorophore. The RNA molecules were then purified using polyacrylamide gel electrophoresis (PAGE) with 8 M urea in a 20% (w/v) acrylamide: bis-acrylamide gel, followed by extraction into buffer (1mM EDTA, 10% v/v acid phenol, 300mM sodium acetate pH 5.0). RNAs were then isolated by ethanol precipitation and dissolved in nuclease-free water (Ambion). RNA products were then quantified by UV-Vis spectroscopy.

3’ phosphate modified U6 molecules for CoSMoS assays were prepared using a double- splinted ligation reaction with T4 RNA ligase 2 and three RNA fragments corresponding to U6 nt 1-12 (synthetic and containing a 5’ biotin and internal Cy5, purchased from IDT), nt 13-94 (*in vitro* transcribed), and nt 95-112 (synthetic, purchased from IDT) and two DNA splints. In the ligation reaction, the three pieces were used at a ratio of 1 (nt 1-12):2 (nt 13-94):2 (nt 95-112):1.5 (DNA splints). Ligation products were gel purified, extracted, isolated, and quantified as above.

### Electrophoretic Mobility Shift Assays (EMSA)

RNAs were first diluted into RNA dilution buffer (100 mM KCl, 20% w/v sucrose, 20 mM HEPES pH 7, 1 mM EDTA, 1 mM TCEP, 0.01% v/v Triton X-100, 0.2 mg/mL yeast tRNA, and 0.2 mg/mL sodium heparin), and proteins were diluted into protein dilution buffer (100 mM KCl, 20% w/v sucrose, 20 mM HEPES pH 7, 1 mM EDTA, 1 mM TCEP, 0.01% v/v Triton X-100, and 0.2 mg/mL BSA) as described (12). Binding was initiated by mixing 5 µL of RNA with 5 µL of protein, and samples were then incubated for 30 min at RT. Mixtures were then loaded onto a pre-run native 6% (w/v) PAGE gel in 1xTBE and separated for 2h at 4°C at 5W. Fluorescence analysis was performed with a Typhoon FLA 9000 scanner (Cytiva) with excitation at 635 nm and results analyzed using ImageJ software (version 1.52v).

### Fluorescence Polarization Assays

Fluorescence polarization experiments were performed according to previous methods (12). In summary, a 2× RNA solution (2 nM, 100 µL) in buffer (100 mM NaCl, 20 mM bis-tris pH 7.0, 10 mM HCl, 1 mM TCEP, 5% v/v glycerol, 0.02 mg yeast tRNA) was combined with 100 µL of protein at various concentrations (0.01-1000 nM) in buffer that also contained 0.2 mg/mL BSA. The mixture was then added to black, non-transparent, flat-bottomed 96-well microplates. Fluorescence polarization data were then collected in triplicate using a Tecan Infinite M1000Pro plate reader with an excitation wavelength of 470 nm and an emission wavelength of 519 nm. The data were normalized to the values obtained with 0 nM protein (the smallest value) and to the highest value, then averaged across three technical replicates. Binding curves were fit using nonlinear regression in GraphPad Prism 4 with a four-parameter logistic equation where FP_min_ and FP_max_ represent the normalized minimum and maximum % bound, *K_D_* stands for the binding dissociation constant, and H represents the Hill coefficient (Eq. 1).

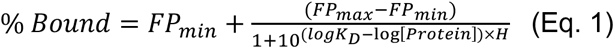

### Microscope Slide Preparation

Microscope slides (100490-396, VWR) and cover slips (12-553-455, Fisher) were cleaned as described (22) using 2% v/v micro-90, 100% ethanol, and 1M KOH with rinsing in between each solution with MilliQ water. After the final rinse, the slides were dried with high purity nitrogen (NI UHP300, Airgas) and aminosilanized with Vectabond (NC9280699, Fisher Scientific). Slides were then passivated by incubation overnight with a mixture of mPEG-SVA (MPEG-SVA-5K, Laysan Bio) and mPEG-biotin-SVA (BIO-PEG-SVA-5K, Laysan Bio) in a 1:100 w/w ratio in 100 mM NaHCO_3_ buffer.

### Single Molecule Microscopy

Single molecule data were obtained using a custom-built objective-type total internal fluorescence microscope system (32). The microscope setup includes a 60× 1.49 NA PlanApo objective (Olympus) and 532 nm (CrystalLaser), 633 nm (Power Technology Inc.), and 785 nm (Power Technology Inc.) lasers for TIRF excitation. Typically, fluorophores were imaged with the 532 nm laser set at a power of 800-900 µW and the 633 nm laser set to 600-700 µW. The 785 nm laser was used for autofocus and was set at 2.5 mW. The emission light was split into <635 nm and >635 nm channels to produce two images that were then imaged on separate sCMOS detectors (Hamamatsu ORCA-Flash4.0 V3) with 2×2-pixel binning. The microscope was controlled using Micro-Manager 2.03 (33).

Prior to data collection, streptavidin-labeled fluorescent beads (T10711, Invitrogen) in PBS were introduced into a channel on the passivated slide to serve as fiduciary markers for alignment. The lane was then washed 0.2 mg/mL streptavidin (SA10-10, Agilent; 50 µL) for 2 min, followed by washing with PBS to remove any unbound beads and streptavidin. Biotin_Cy5_U6 snRNA was diluted to 10 pM in imaging buffer (20 mM HEPES pH 7.0, 100 mM KCl, 1mM EDTA, 20% w/v sucrose, 10 mg/mL BSA, 5 mM protocatechuic acid (PCA), 1U/mL protocatechuate dioxygenase (PCD), 1 mM trolox, 2% v/v DMSO) and incubated in the lane for 1 min, followed by another wash with imaging buffer. A 100 µL solution containing varying concentrations of Prp24, Lsm2-8 and Lhp1 in imaging buffer was added and alternating laser excitation imaging was performed based on the following excitation scheme: The 532 nm and 633 nm lasers were alternatively turned on (1 s/fr) to capture images with an ∼0.5 s lag between each image for ∼1800 s.

### Microscopy Data Analysis

The single molecule data were analyzed using custom MATLAB software (smtoolbox, available at https://github.com/David-Scott-White/cosmos-toolbox). The analysis procedure involved several steps: 1) a mapping file that correlated the positions in the <635 nm channel to the >635 nm channel was created utilizing the fluorescence beads as fiducial markers, 2) lateral drift was corrected by computing a non-reflective similarity transformation between temporally separated images, 3) potential areas of interest (AOIs) containing single RNA molecules were identified in the >635 nm channel by averaging the first five frames and applying a generalized likelihood ratio test to the resulting image. These detected AOIs were then fit to a two-dimensional Gaussian within a 5×5 pixel area, 4) the accepted AOIs were subsequently mapped to the <635 nm channel using the mapping transformations, 5) the fluorescence trajectories in the <635 nm channel (Lsm2-8 proteins) for each AOI were processed using the divisive segmentation and clustering algorithm (DISC, (34)) to interpret each trajectory as a binary signal indicating bound or unbound states over time.

The durations of bound and unbound events (dwells) were treated as single or double exponential distributions, and the underlying parameters of each distribution were estimated using maximum likelihood methods (35, 36). A comparison was made between models described by single exponential terms, the sum of multiple exponential terms, or gamma distributions. The log likelihood-ratio (LLR) test was employed for model selection. For exponential distributions, the likelihood of observing a dwell time of a specific duration is expressed as (Eq. 1):

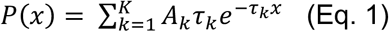

where the variable *k* denotes the number of exponentials being estimated, *A* represents the amplitude associated with a particular exponential term, and 𝜏 signifies the estimated time constant corresponding to that term.

To control photobleaching, a biotinylated SNAP protein was derivatized with SNAP- Surface® 549, immobilized on a slide, and its fluorescence lifetime measured under identical conditions as previously described (22).

### Kinetic Analysis Using QuB

To analyze the binding kinetics of Lsm2-8 to U6 RNA, we performed Hidden Markov Model (HMM) fitting in QuB using single-molecule trajectories collected at 1 nM, 3 nM, and 10 nM Lsm2- 8 concentrations (37). Transition rates were optimized using Maximum Idealized Point (MIP) likelihood rate estimation, which globally optimizes state transition probabilities across all observed molecules(38). A range of user-defined models with different numbers of states (two to four) and transition pathways were tested to identify the best kinetic description of Lsm2-8 binding. Model selection was based on the Bayesian Information Criterion (BIC) (39):

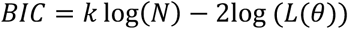

where *k* is the number of free parameters, 𝑁 is the number of frames analyzed, and 𝐿(θ) is the log-likelihood of the dataset given the model. After evaluating multiple models, we found that a model with six free parameters provided the best fit, as indicated by the lowest BIC value. The final kinetic scheme, presented in **Fig. S4**, represents the key states and transition pathways identified from the optimal model.

## DATA AVAILABILITY

Raw microscopy data and gel images have been uploaded to Zenodo DOI: 10.5281/zenodo.14790398 and will be released upon manuscript acceptance.

## SUPPORTING INFORMATION

This manuscript contains supporting information, including supplementary figures illustrating protein and RNA purification results, spot accumulation curves, photobleaching controls, and kinetic models. It also includes data tables showing fitted binding affinity parameters, bound and unbound time distributions, and oligonucleotide sequences. These materials can be accessed online alongside the full text of the article.

## ACKNOWLEDGEMENTS

We thank Dr. David White for help with data analysis protocols and code, and Dr. Harpreet Kaur for assistance with Lsm2-8 purification.

## FUNDING AND ADDITIONAL INFORMATION

This work was supported by grants from the National Institutes of Health (R35 GM136261 to AAH and R35 GM118131 to SEB).

