## Supplemental Figures and Tables for "RNA Modifications and Prp24 Coordinate Lsm2-8 Binding Dynamics during *S. cerevisiae* U6 snRNP Assembly"

Ye Liu[1], Yuichiro Nomura[1], Samuel E. Butcher[1] and Aaron A. Hoskins[1,2,*]

[1] Department of Biochemistry, University of Wisconsin-Madison, Madison, WI 53706

[2] Department of Chemistry, University of Wisconsin-Madison, Madison, WI 53706

This supplementary file contains:

Supplementary Figures S1-S4

Supplementary Tables S1-S4

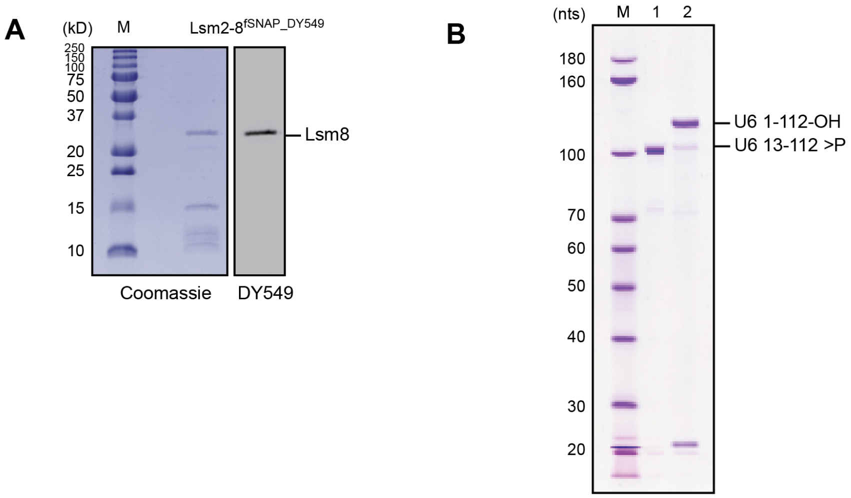


**Supplementary Fig S1. Denaturing PAGE analysis of fluorophore-labelled reagents. (A)** SDS-PAGE analysis of the purified Lsm2-8$^{fSNAP}$ labeled with DY549. **(B)** Denaturing urea PAGE gel showing the U6 13-112 transcripts (lane 1) and final U6 1-112 ligation product (lane 2).

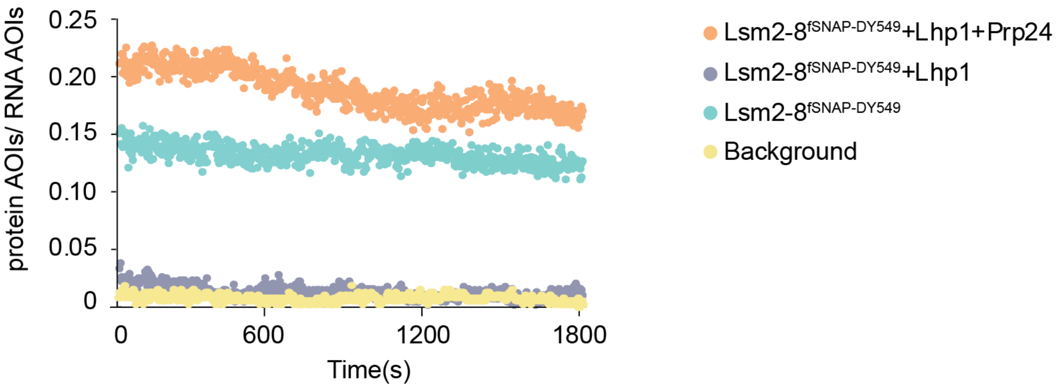


**Supplementary Figure S2. Surface accumulation of fluorescent Lsm2-8$^{fSNAP}$ molecules.** Spot accumulation trajectories (normalized to the number of immobilized RNAs or blank AOIs (background)) are compared for the Lsm2-8 under conditions in which the U6 RNA 3' end is freely accessible (teal, orange) or bound by Lhp1 (purple).

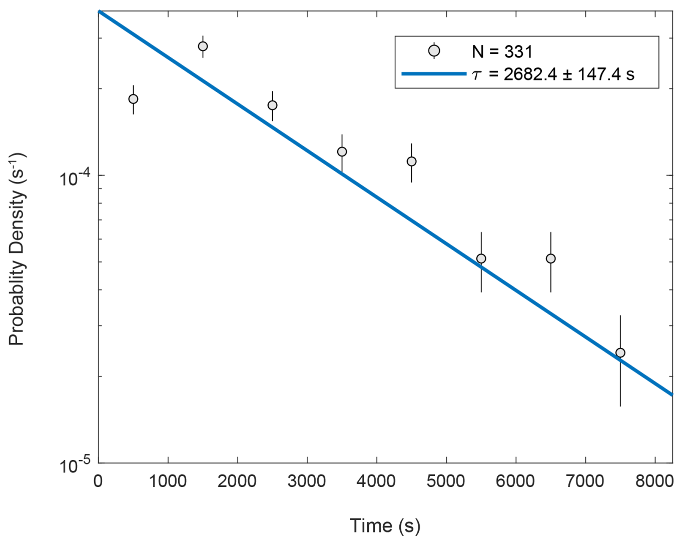


**Supplementary Figure S3. Fluorescent lifetime of DY549-labelled fSNAP protein.** Biotinylated fSNAP protein labeled with DY549 was immobilized on a glass slide, and imaged under the same conditions as the CoSMoS assays used for monitoring Lsm2-8 dynamics. The lifetime distribution was fitted to a single exponential equation, yielding a time constant ($\tau$) of 2682.4±147.5s.

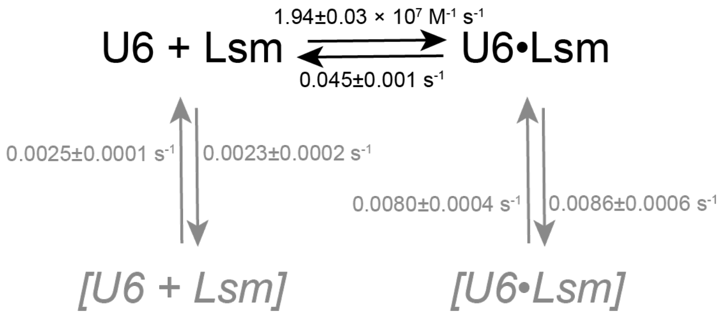


**Supplementary Figure S4. A kinetic model for the interaction between 3' end modified U6 RNA and the Lsm2-8 complex.** The 3' end modified U6 snRNA (U6; 1-112-P) can reversibly bind with the Lsm2-8 complex (Lsm). At slower rates, the substrates and products can occasionally transition to states from which the RNA and protein can neither directly associate or dissociate (grey arrows and italicized complexes).

**Supplementary Table 1.** Fitted binding affinity parameters from fluorescence polarization assays.

| RNA | Lsm2-8$^{WT}$ | | | Lsm2-8$^{fSNAP}$ | | |
|---|---|---|---|---|---|---|
| **U6 3' tail** | **$K_D$ (nM)** | **Hill coefficient** | **$R^2$** | **$K_d$ (nM)** | **Hill coefficient** | **$R^2$** |
| U6 95-113-OH | 6.75±1.15 | 1.18±0.24 | 0.98 | 6.70±1.10 | 1.12±0.22 | 0.98 |
| U6 95-112-OH | 84.64±16.82 | 1.26±0.26 | 0.97 | 113.80±16.80 | 1.35±0.24 | 0.98 |
| U6 95-112-P | 10.34±4.10 | 1.11±0.45 | 0.89 | 12.83±3.87 | 0.88±0.23 | 0.95 |

**Supplementary Table 2. Fitted unbound time kinetic parameters from dwell time analysis.**

| Tagged RNA | Other protein added | Amplitude 1 | T1 (s) | Amplitude 2 | T2 (s) | Number of Events Fit |
|---|---|---|---|---|---|---|
| U6 1-113-OH | - | 0.50±0.03 | 85.60±10.40 | 0.50±0.03 | 559.80±112.30 | 997 |
| U6 1-112-P | - | 0.81±0.03 | 32.40±6.40 | 0.19±0.03 | 370.75±28.45 | 2401 |
| U6 1-112-OH | - | 0.72±0.06 | 166.05±82.95 | 0.28±0.06 | 333.05±52.95 | 2657 |
| U6 1-113+5U-OH | - | 0.67±0.01 | 117.70±3.25 | 0.33±0.01 | 403.50±29.85 | 1113 |
| U6 1-113-OH | Prp24 | 0.60±0.04 | 33.35±16.65 | 0.40±0.04 | 354.15±51.25 | 1142 |
| U6 1-112-P | Prp24 | 0.65±0.03 | 42.60±12.00 | 0.36±0.03 | 404.45±146.45 | 2303 |
| U6 1-112-OH | Prp24 | 0.83±0.02 | 74.10±8.00 | 0.17±0.02 | 280.10±69.9 | 1461 |
| U6 1-113+5U-OH | Prp24 | 0.75±0.03 | 120.40±14.90 | 0.25±0.03 | 382.20±24.10 | 2827 |
| U6 1-112-P | $Prp24^{\Delta SNFFL}$ | 0.82±0.07 | 112.80±8.30 | 0.18±0.07 | 442.90±38.70 | 1668 |
| U6 1-113+5U-OH | $Prp24^{\Delta SNFFL}$ | 0.70±0.03 | 126.80±9.10 | 0.30±0.03 | 399.30±4.20 | 1991 |

**Supplementary Table 3. Fitted bound time kinetic parameters from dwell time analysis.**

| Tagged RNA | Other protein added | Amplitude 1 | T1 (s) | Amplitude 2 (s) | T2 | Number of Events Fit |
|---|---|---|---|---|---|---|
| U6 1-113-OH | - | 0.80±0.10 | 20.85±8.75 | 0.20±0.10 | 183.10±8.60 | 911 |
| U6 1-112-P | - | 0.79±0.07 | 30.90±14.90 | 0.21±0.07 | 226.80±54.30 | 2402 |
| U6 1-112-OH | - | 0.99±0.01 | 12.40±3.60 | 0.01±0.01 | 304.60±199.50 | 2454 |
| U6 1-113+5U-OH | - | 0.88±0.02 | 20.90±1.80 | 0.12±0.02 | 194.60±80.10 | 1078 |
| U6 1-113-OH | Prp24 | 0.45±0.03 | 29.40±6.10 | 0.55±0.03 | 290.30±58.50 | 1121 |
| U6 1-112-P | Prp24 | 0.69±0.06 | 32.4±4.6 | 0.31±0.06 | 295.40±160.60 | 2337 |
| U6 1-112-OH | Prp24 | 0.88±0.07 | 19.70±1.30 | 0.12±0.07 | 282.20±48.90 | 1406 |
| U6 1-113+5U-OH | Prp24 | 0.90±0.03 | 24.80±1.00 | 0.10±0.03 | 255.90±5.50 | 2642 |
| U6 1-112-P | Prp24$^{\Delta SNFFL}$ | 0.85±0.01 | 19.70±0.70 | 0.15±0.01 | 143.25±24.45 | 1519 |
| U6 1-113+5U-OH | Prp24$^{\Delta SNFFL}$ | 0.89±0.01 | 22.60±1.70 | 0.21±0.01 | 267.10±72.50 | 1802 |

**Supplementary Table 4. Synthetic Oligonucleotide Sequences.**

| Oligo # | ID | Sequence (5'-3') | Length (nts) |
|---|---|---|---|
| 1 | FAM_U6_95-113-OH | 5' - /56-FAM/ rArGrArGrArUrUrUrArUrU rUrCrGrUrUrUrUrU - 3' | 19 |
| 2 | FAM_U6_95-112-P | 5' - /56-FAM/ rArGrArGrArUrUrUrArUrU rUrCrGrUrUrUrU/3Phos/ - 3' | 18 |
| 3 | FAM_U6_95-112-OH | 5' - /56-FAM/ rArGrArGrArUrUrUrArUrU rUrCrGrUrUrUrU - 3' | 18 |
| 4 | Bio_Cy5_U6_1-12 | 5' - /5Biosg/ rGrUrUrCrGrC/iCy5/rGrAr ArGrUrA- 3' | 12 |
| 5 | DNA 12-13 bridge | 5' - GTCCACGAAGGGTTAC TTCGCGAAC - 3' | 25 |
| 6 | U6 95-112-P | 5' - /5Phos/rArGrArGrArUrUr UrArUrUrUrCrGrUrUrUrU /3Phos/ - 3' | 18 |
| 7 | DNA 94-95 bridge | 5' – GAAATAAATCTCTTTGT AAAACGGTTC - 3' | 27 |
